# A Transformer based method for the Cap Analysis of Gene Expression and Gene Expression Tag associated 5’ cap site prediction in RNA

**DOI:** 10.1101/2025.09.21.677558

**Authors:** Dibya Kanti Haldar, Avik Pramanick, Chandrama Mukherjee, Pralay Mitra

**Affiliations:** Centre for Computational and Data Sciences, Indian Institute of Technology Kharagpur, West Bengal, India-721302; Department of Computer Science and Engineering, Indian Institute of Technology Kharagpur, West Bengal, India-721302; Institute of Health Sciences, Presidency University, Kolkata, West Bengal, India - 700073

**Keywords:** Cap Analysis of Gene Expression, Capping, Transcription Start Site, Transformer, bi-LSTM

## Abstract

5’ RNA capping is one of the major post-transcriptional modifications for the mobility and stability of RNA molecules. Measuring 5’ caps of RNAs can help quantify expression levels of mRNAs and lncRNAs. One of the most successful RNAseq methods that have used capping as a tool to quantify expression of transcription is Cap Analysis of Gene Expression(CAGE). Computational prediction of capping can therefore be used as a precursor to the prediction of transcriptional expression. Unfortunately, there is hardly any computational technique that has focused purely on predicting 5’ capping. We have developed a transformer-based method for computational prediction of capping from DNA sequences.

Our Llama and ReLoRA-based pre-training model, and Llama and LoRA-based fine-tuning model predict 5’ cap sites. We have used Leave-one-chromosome-out-cross-validation for our model. The average accuracy, and F1-score after fine-tuning the human genome hg19(mouse genome mm9) for sequence classification is 79.12%(78.09%), and 78.11%(76.17%), respectively. We noted attention peak-based motifs having an aggregate Wilcoxon rank-sum p-value of 1.075e-10 between the attention peak region and the entire context window for the predicted positive motifs; an aggregate p-value of 7.17e-18 for the predicted negative motifs; and an aggregate p-value of 6.70e-08 between the attention peaks of the predicted positive and the predicted negative motifs. Our Llama-based approach aims to create a sequence-based framework to identify 5’ capping sites corresponding to CAGE peaks. Our analysis reveals statistically significant motifs from the regions of peak attention scores, which demonstrates biological relevance for some through their resident sites matching with known TF motifs.

## 1. Introduction

The eukaryotic cell, biological unit of all multi-cellular life, has three components encapsulating all the information and carrying out all major activities, DNA, RNA and proteins. In the process, the pre mRNAs undergo post-transcriptional modifications to become RNA, and synthesize proteins. The post-transcriptional modifications are poly-adenylation, capping, methylation and phosphorylation.

Capping (Pelletier et al., 2021; Ramanathan et al., 2016) is the process of the addition of a 7 methyl-guanosine cap to the 5’ end of all RNA polymerase transcribed RNAs, after which the capped RNA can cross the nuclear membrane and traverse through the cytoplasm for translation. Being a post-transcriptional modification, RNA 5’ capping is essential for the translation initiation, and the prevention of the exonuclease-based degradation of the RNA 5’ region.(Potužník & Cahova, 2024) Decapping of RNA is believed to be destined for degradation. However, this concept is challenged after the identification of cytoplasmic recapping of a subset of mRNAs as well as lncRNAs.(Borden et al., 2021; C. Mukherjee et al., 2012) Recapping can occur at the sites other than Transcription start site (TSS), upstream of the promoter, as evidenced by the capping of decaying RNA and the capping of spliced exons.(Fejes-Toth et al., 2009; Trotman & Schoenberg, 2019) Therefore, the cap trapping protocol would also identify caps at sites downstream of TSS. RNA capping is a regulatable activity, and the status of RNA capping is not fixed. It can fluctuate during different cellular conditions including but not limited to differentiation and development. Well known methods to identify cap structures include CapIP, CapTure, CapQ, Oligocapping, CapSMART, CapTrapper, CapQuant, and CAGE.(Borden et al., 2021) New cap structures i.e. those not based on the 7 methyl-guanosine cap, were identified in human and mouse cells using CapQuant8, and the identified caps are based on alternate enzymes and mechanisms like FAD, NAD+, NADH, UDP-Glc, UDP-GlcNAc, mGpppmA.(Bird et al., 2016; Wang et al., 2019; Wiedermannová et al., 2021) CAGE or cap analysis of gene expression (Kodzius et al., 2006; Shiraki et al., 2003) is a popular method of RNA sequencing, originally performed by counting the RNA 5’ caps, captured using the cap trapping protocol.(Takahashi et al., 2011) CAGE is used primarily for TSS annotation, along with methods like RAMPAGE, GRO-cap, STRT, NanoCAGE, etc. (Adiconis et al., 2018) Even so, CAGE analysis has identified that around 25% of the capping sites in mammalian genome were located at non promoter regions, namely at spliced exons (Kiss et al., 2015)

Conversely, CAGE has only a recall rate of 75% in TSS peak identification.(Adiconis et al., 2018) Therefore, capping occurs near promoter sites as well as some sites across the gene, making capping prediction a problem disparate from TSS prediction. The first CAGE protocol involved cap trapping at the 5’ ends, synthesis of cDNA fragments using oligo-dT primers, tag cleaving using restriction enzymes, followed by Sanger (Crossley et al., 2020) sequencing. DeepCAGE (de Hoon & Hayashizaki, 2008) added barcode multiplexing to the protocol. Subsequent methods like CAGEscan (Bertin et al., 2017; Plessy et al., 2010) and HeliScopeCAGE (Itoh et al., 2012; Kanamori-Katayama et al., 2011) skipped the enzymatic tag cleavage part of the original protocol. The CAGE protocol generates CAGE tags for each of the sequence fragments it analyzes, and the CAGE peaks are derived from the overlapping tags. The CAGE protocol can identify TSS with a very high precision, and sometimes the CAGE peaks can be down to a single nucleotide, suggesting a single nucleotide precision. CAGE peaks usually range from 1 to 50 bps, and are usually found in the first few hundred kilobases from TSS signals. CAGE peaks frequently overlap with H3K4me3 peaks and DNaseseq peaks.(Adiconis et al., 2018)

The TSS can be signalled by motifs like the TATA box, the Initiator element, the promoter upstream element, and the promoter downstream element. The presence of TSS signals do not by themselves indicate the start of transcription, but a possibility of one. Sometimes TSS signals are present in isolation, and are referred to as sharp TSS signals. Other times, they might be present in a cluster of overlapping or nearby TSS signals, referred to as a broad TSS signal. The TATA box is one of the most prominent TSS signals, and is represented by the pattern TATAWADR i.e. TATA[A/T]A[G/A/T][G/A]. It is recognized by the Tata Binding Protein (TBP), a general Transcription Factor, and associated with TBP Associated Factors (TAFs).(Mishal & Luna-Arias, 2022) The Initiator is also a common TSS motif, and is identified by the pattern YYANWYY i.e. [T/C][T/C]A[A/T/G/C][A/T][T/C][T/C]. The Promoter Upstream Element, usually located upstream of the TATA box motif, is recognized by the pattern SSRCGCC ([G/C][G/C][G/A]CGCC), whereas the Promoter Downstream Element, usually located downstream of the Initiator, is identified by the pattern RGWYV i.e. [G/A]G[A/T] [T/C][G/C/A].

Differential gene expression is the mechanism responsible for pluripotency in the Eukaryotic organisms. Gene expression is dependent on factors such as the presence of cell type specific TFs, chromatin accessibility of the gene, methylation state of the underlying histones and of the gene itself, the active enhancers and silencers regulating the gene, the transcription activation and repression mechanisms, regulatory non-coding RNAs. This makes the quantification of gene expression a multifactor problem, many of which can be obtained from the DNA sequence itself, like enhancer and silencer sites. Other factors, like associated TFs, regulatory RNAs, and methylation state of the underlying histones, can be identified using other mechanisms like protein expression for TFs expressed in cell/-tissue, gene regulatory networks for regulatory RNAs and ChIP seq for analyzing histone modifications. While this makes it difficult to identify the active TSS from sequence alone, there have been attempts to identify them from sequence alone.

Capping sites, on the other hand, can be identified from DNA sequence alone, even though attempts to predict capping sites have been scarce. While the CAGE protocol maps the capped RNA, the CAGE peaks are often clustered and not found in the same sites consistently, with the average deviation between nearby CAGE peaks being 51 nucle-otides; reducing the reliability of the CAGE peaks as narrow capping sites. Increasing the context length to incorporate more DNA around CAGE peaks, would allow us to reliably predict broad capping sites. However, CAGE peak sites are not found to be associated with any known motif, and TSS motifs like promoters and initiators are usually present at a variable distance from the CAGE peaks. In large context of 500 or 1000 bps, sequences in possession of CAGE peaks as well as sequences without, have been found to possess similar number of TSS signals in them.

We have developed a framework that takes the DNA subsequence as input and outputs the predicted capping sites. Our framework, first divides the input DNA sequence into words of size 8 nucleotides where each word shares the first 4 nucleotides with the previous and the next 4 nucleotides with the next. We repeated this process with offsets of 1, 2 and 3, before passing it through a Byte Pair Encoding based Tokenizer, and to prevent motifs from being missed out. The tokens are then passed through our Llama (Touvron et al., 2023) and ReLoRA (Lialin et al., 2023) based pre-training module using a learning rate scheduler. The pretraining is followed by a Llama and LoRa (Hu et al., 2022) based finetuning. We have trained our model using Leave One Chromosome Out Cross Validation (LOCOCV) (Tahir et al., 2025), using a 1:1 proportion of positive and negative samples, in order to prevent data leakage and conform to standard dataset creation practices. Our model achieved an LOCOCV capping classification accuracy of 79.12±1.16% (78.25±1.00%), a precision of 78.64±2.10% (79.84±3.50%), and a recall of 77.66±2.11% (73.49±3.275%), along with an F1 score of 78.11±1.13% (76.40±1.01%) on the human (mouse) genome using only Llama and LoRa based fine-tuning on our pretrained model for a context length of 512.

## 2. Materials and Methods

### 2.1 Dataset details

We created deep learning techniques for 5’ cap detection by training the models on the human genome hg19 and mouse genome mm9. The FANTOM5 (Noguchi et al., 2017) consortium have bed files containing CAGE peaks for the human genome hg19 and the mouse genome mm9. The CAGE peaks represented the genomic locations of the capping sites found through CAGE. We downloaded the human genome hg19 and the mouse genome mm9. For creating the dataset, we downloaded the hg19 and mm9 fasta sequences adjacent to the CAGE peaks, downloaded from FANTOM5, which included a near equal number of nucleotides upstream and downstream to the peak. For our site prediction task, we divide the genomic sequence into fixed size subsequences and then apply binary classification on the subsequences. Positive sequences were those which contained CAGE peaks, whereas negative sequences had the CAGE peaks at a considerable distance away from them. The number of positive and negative sequences obtained from the human genome hg19 and the mouse genome mm9 are given below in Table 1.

**Table 1:** Chromosomes and corresponding number of positive and negative samples (hg19 and mm9). The hg19 positive sequences return the number of samples per chromosome in our dataset having CAGE peaks taken from the hg19 human genome. The mm9 positive sequences return the same from the mm9 mouse genome. The hg19 negative sequences and mm9 negative sequences return the number of samples per chromosome in the dataset not having CAGE peaks taken from hg19 and mm9 genomes respectively.

| chr | Human<br>positive | (hg19)<br>negative | Mouse<br>positive | (mm9)<br>negative | chr | Human<br>positive | (hg19)<br>negative | Mouse<br>positive | (mm9)<br>negative |
| --- | --- | --- | --- | --- | --- | --- | --- | --- | --- |
| **1** | 18654 | 20923 | 10089 | 11284 | **13** | 4145 | 4783 | 6005 | 6750 |
| **2** | 16437 | 18028 | 12369 | 13580 | **14** | 7031 | 7586 | 6201 | 6769 |
| **3** | 11514 | 12845 | 8069 | 8858 | **15** | 6417 | 7245 | 6439 | 7046 |
| **4** | 896 | 10035 | 8714 | 9721 | **16** | 6832 | 7390 | 5531 | 6115 |
| **5** | 9953 | 11169 | 9645 | 10918 | **17** | 10370 | 11323 | 7537 | 8241 |
| **6** | 11479 | 12518 | 9488 | 10571 | **18** | 3648 | 4295 | 4278 | 4948 |
| **7** | 10410 | 11700 | 11803 | 13057 | **19** | 10315 | 11124 | 5938 | 6523 |
| **8** | 8608 | 9698 | 7536 | 8654 | **20** | 5483 | 6275 | 0 | 0 |
| **9** | 7669 | 8396 | 8641 | 9330 | **21** | 2563 | 2778 | 0 | 0 |
| **10** | 8747 | 9672 | 7789 | 8631 | **22** | 3873 | 4023 | 0 | 0 |
| **11** | 11149 | 12282 | 12475 | 13744 | **X** | 5789 | 6453 | 4303 | 4819 |
| **12** | 10993 | 12418 | 5721 | 6557 | **Y** | 207 | 218 | 32 | 29 |

We constructed the positive samples by taking the DNA segments containing the CAGE peaks, as well as 256 nucleotides upstream and downstream of the CAGE peak start sites. We built the negative samples by taking 512 nucleotides long DNA segments randomly between 10000 bps and 20000 bps upstream of the CAGE peak start sites, while taking care that these segments do not overlap with the DNA segments in the positive samples. To avoid any misinterpretation, we had to remove those positive samples whose DNA segments had overlapping base pairs with some negative sample, resulting in approximately 10% greater number of negative samples per chromosome than the corresponding positive ones.

Our pretraining dataset was created by combining the genomes of Homo Sapiens (human), Mus Musculus (mouse), Pan Troglodytes (chimpanzee), and Pan Paniscus (bonobo), and dividing them into fragments of size 512 for a context window of size 512.

### 2.2 Method

Our approach, as shown in the flow diagram in Figure 1 used Eleuther AI’s GPTNeoX-20B tokenizer, a tokenizer trained on the diverse and extensive PILE (Gao et al., 2020) dataset, which is based on Byte Pair Encoding based sub-word tokenization. Byte Pair Encoding has characteristics similar to the merge sort algorithm, and starts the tokenization process from tokens of size 1, in other words, character level tokens. The most frequent token pair from all pairs of adjacent tokens is taken and concatenated into a single token. This is repeated for *k* number of steps, and ends up building the vocabulary. This tokenization is able to obtain most of the prefixes, stem words and postfixes as tokens when trained on the English text corpus.

**Figure 1:**
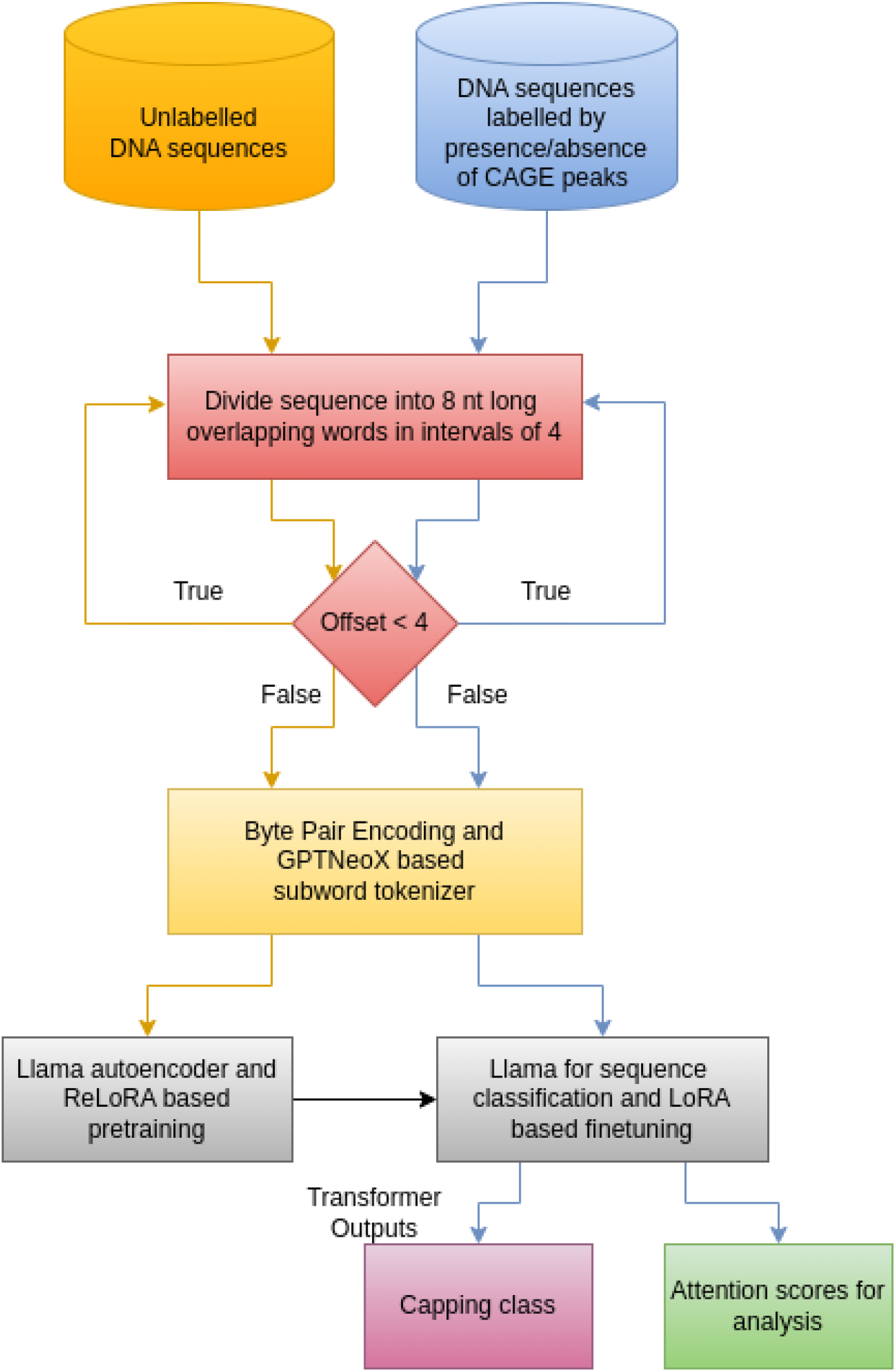
Flow diagram of our approach, showing DNA sequences divided into words, tokenized using Byte Pair Encoding based tokenizer, pretrained (orange arrows) using Llama and ReLoRA based pretraining, and finetuned (blue arrows) using Llama and LoRA based finetuning. Please note the processes remain same, pretraining and finetuning varies in input and output.

We divide DNA sequences of size 512 nucleotides into overlapping tokens of size 8. Therefore, each token, except for the first and the last, shares its first 4 nucleotides with the previous token and its last 4 nucleotides with the next token. We repeat the tokenization for offsets of 1, 2 and 3, and append them to the list of tokens. The tokens are then encoded with the GPTNeoX-20B tokenizer, creating a vocab of size 50254.

Next, we pretrained our pretraining dataset on base Llama 20m for 1000 steps, before switching to ReLoRA optimized Llama 20m for a further 13197 steps. The Llama model had 28.17 million parameters and comprised of a positional embedding layer of size 50254×256, 4 heads and thus a 4*x* llama decoder layer, each comprised of 4 projections: query, key, value and output, followed by Llama Rotary Embedding, and a Llama MLP comprised of 3 projections: gate projection, down projection, and up projection.

LoRA factorizes weight updates into low rank matrices, allowing the model to use fewer total updates, and thus lower trainable parameters, during fine-tuning. ReLoRA extends this mechanism to the pre-training step, allowing the complete model to be trained using less GPU memory after training the full model for a few warm-up steps. Consequently, ReLoRA linear layer replaced all the linear projection layers in our llama-based model.

The training loss and the validation loss fell continuously in lockstep, with the training loss reaching 0.25195, down from its peak at 11.0 at the beginning, and the validation loss reaching 0.22348, down from its peak at 5.36977 at step 200, when it was first recorded. Only the model tokenized using offsets was able to break below the loss of 1.00, whereas the models tokenized without using offsets failed to do so.

We followed this by tokenizing our training and validation datasets, which would be used during fine-tuning. We tokenized our training and validation datasets using the same tokenizer as during pre-training, having the same vocab of 50254 tokens. Our datasets were fine-tuned using llama for sequence classification, involving 16 million parameters model for 5 epochs and 23440 steps. The fine-tuning used LoRA and consisted of 4 attention heads, each comprising of self-attention with LoRA query, key, value, and output projections, and 4 hidden layers of LoRA MLP down projection, gate projection, and up projection.

## 3. Results

### 3.1 Training and Validation

Our Llama and ReLora based pre-trained and fine-tuned capping prediction model was trained using LOCOCV on the human genome hg19. It was pre-trained on DNA sequences having a context length of 512, using a Llama transformer having 20 million parameters. It was trained on the original Llama model for 1000 steps, and further trained on the ReLoRA based Llama model for another 13197 steps. After fine-tuning the human genome hg19 on llama for sequence classification for 5 epochs on a batch size of 64 for 23440 steps, we obtained an accuracy of 79.12±1.16%, a precision of 78.64±2.10%, a re-call of 77.66±2.11%, and an F1 score of 78.11±1.11% on 24 chromosomes of the human genome using LOCOCV (Figure S1). Similarly, after fine-tuning the mouse genome mm9 on llama for sequence classification for 5 epochs on a batch size of 64 for 23075 steps, we obtained an LOCOCV accuracy of 78.09±1.08%, a precision of 79.78±3.32%, a recall of 73.105±3.36%, and an F1 score of 76.17±1.23% (Figure S2).

A bigger context window allows us to find more motifs in the promoter neighborhood, such as the Promoter Downstream Element, Promoter Upstream Elements, silencers, and enhancers. A smaller context window means access to less information about the neighborhood of the capping sites. The capping predictions are validated using the CAGE peaks, which though not the exact capping sites, usually are shifted by some variable number of nucleotides, since the CAGE protocol measures the CAGE tags in bulk, and the capping sites of different cells usually do not agree. The returned CAGE tags are not located near known TSS signals like TATA boxes and Initiators, and can be found at various distances from the TSS signals, with the average distance being around 260 for TATA boxes and 91 for initiator signals. The CAGE peaks also keep shifting and are not found in a common location, since we have found multiple CAGE peaks in the same genomic neighborhood.

We also searched for differentiating capping motifs by comparing the vocab generated by the attention peaks of both predicted classes, and we obtained a few of them. Since the TSS motifs like TATA box, initiator and the Promoter Downstream Element typically contains 8 nucleotides, we too compared the vocabs using nucloetide motifs of length 8. We compared the distribution of these motifs between the entire context window and the attention peak window. Using the Mann-Whitney U test, also known as the Wilcoxon rank-sum test, we obtained the p-value of the normalized attention peak motif frequencies against the normalized motif frequencies in the full context window for the predicted positive samples as 1.0753e-10. For the predicted negative samples, we got the p-value as 7.1697e-18. We also compared the motif frequencies between the attention peaks of the predicted positive samples and the predicted negative samples, and obtained the p-value as 6.6977e-08.

After normalizing by the ratio of total vocab frequencies in the positive and the negative classes, we computed the motif-wise p-values of the predicted positive samples by comparing the token count vectors of the predicted positive samples with that of the predicted negative samples. In an order of increasing p-value, the top 10 motifs were TCTTGAAT, GTCTTGAA, ACTCATGT, CACTCATG, CCCTCAGC, TGATCTGC, TTTTTATT, CAATACAT, GATGTGAG, TTGATCTG. These predicted motifs along with their p-values, and class-wise frequencies, found in the regions having the highest attention scores, are shown in Figure 2. There are 6 more predicted motifs with p-values less than 0.01, namely AATACATA, ATTTAGGG, TAATTTCA, CTTGAATG, ATTCATAG, and ATGTGAGT.

**Figure 2:**
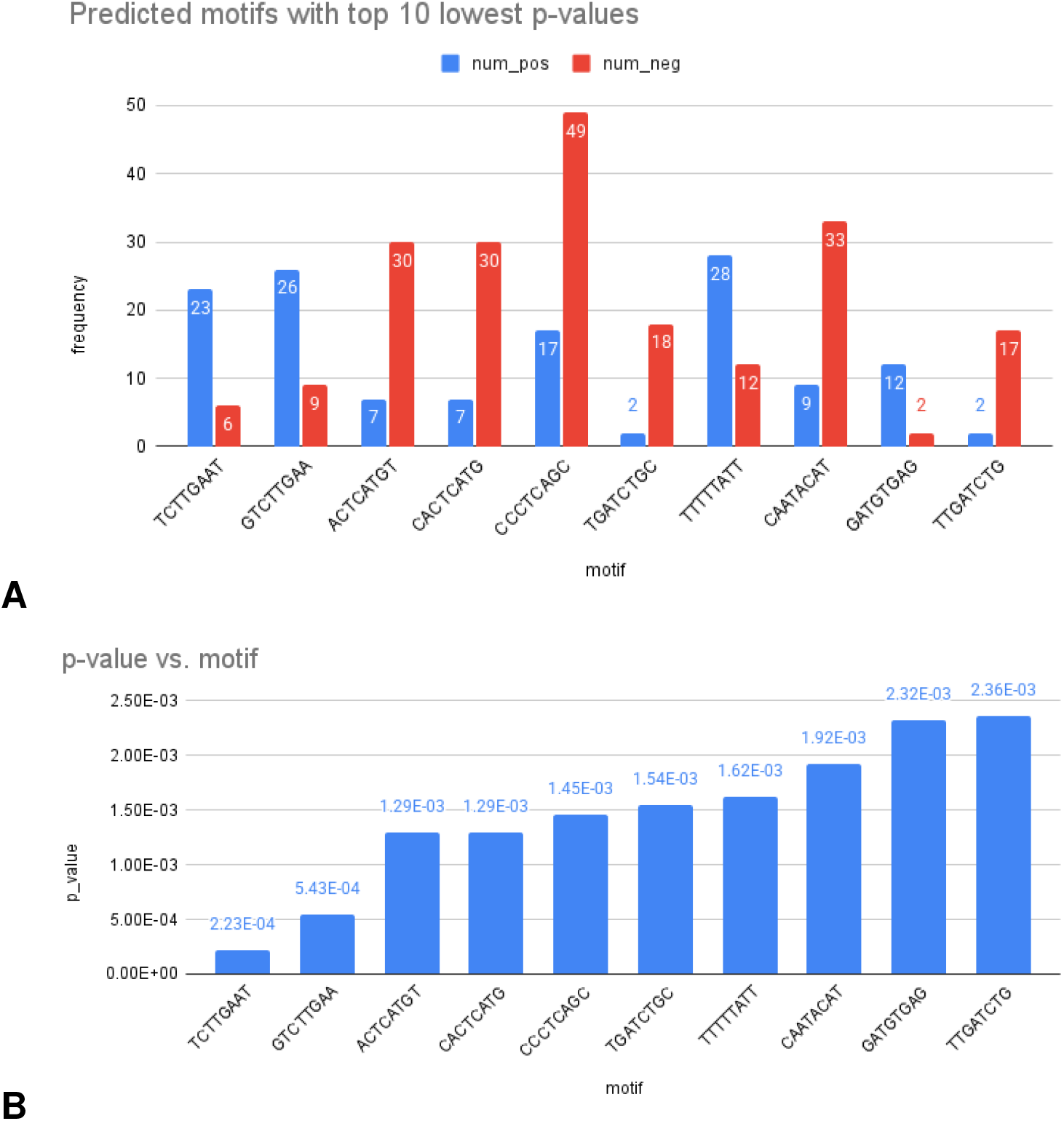
A chart showing A) predicted motifs with the top 10 lowest p-values and B) their respective frequencies in regions of peak attention scores for positive class(blue) and negative class(red)

One of the above motifs TTTTTATT is found to match with the binding sites of the TF motifs MA0497.1, MA0773.1, MA0846.1, MA0846.1, and MA0052.4 using only the attention token ± 5 nts. TTTTTTATT is associated with MA0497.1 with an FDR of 18.43%, a p-value of 3.79e-5 and a q-value of 1.42e-4. It is associated with MA0773.1 with an FDR of 13.95%, a p-value of 6.52e-5 and a q-value of 2.18e-4. It is associated with the TF motif MA0846.1 with an FDR of 24.56%, with 3 matches all having p-values of 2.2e-5 and q-values of 3.03e-4, and 6 matches all having p-values of 7.78e-5 and q-values of 3.57e-4. It is also associated with the TF motif MA0052.4 with an FDR of 15.68%, a p-value of 2.26e-5 a and a q-value of 7.24e-5. When looking at the predicted sequence motifs using the attention token ± 10 nts, we get ATGTGAGT with a prediction based p-value of 0.00786. It matches the binding site of the TF motifs MA0601.1, MA0135.1 and MA0791.1. It is associated with MA0601.1 with an FDR of 41.58%, a p-value of 8.95e-6 and a q-value of 4.65e-4. It’s associated with MA0135.1 with an FDR of 43.58%, and a q-value of 5e-4. For 2 matches, the p-values are 7.95e-6 and for the other 12 matches, the p-values are all 1.53e-5. Its also associated with the TF motif MA0791.1 with an FDR of 48.37%. For 10 of the matches, the p-values are all 4.1 e-7 and the q-values are all 1.9e-5. For the 2 remaining matches, the p-values are 1.96e-6 and the q-values are 5.09e-5. These associations showcase the statistical significance of the predicted motifs. The process to find these motifs is elaborated in Figure 3

**Figure 3:**
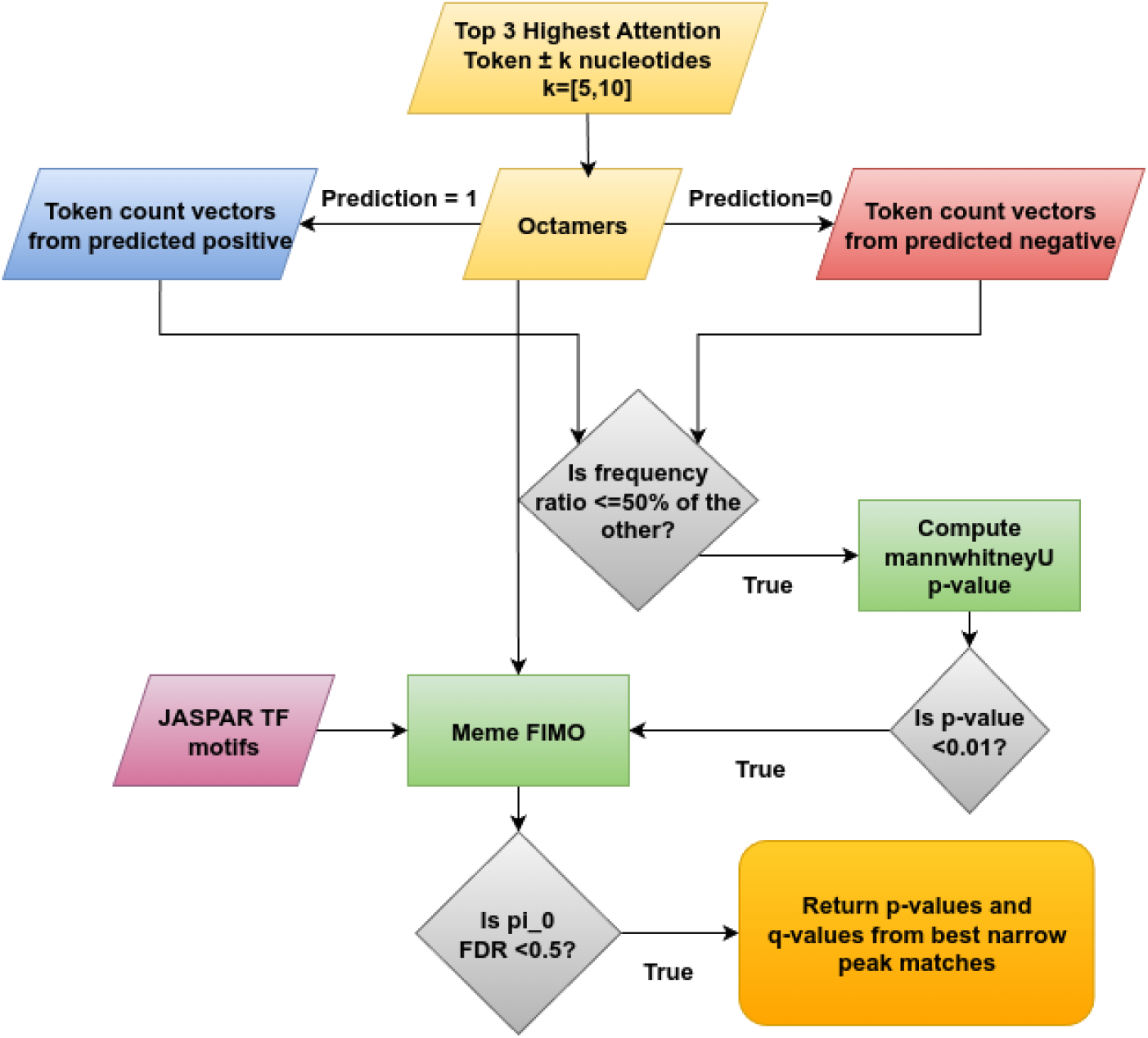
Flow diagram of the process of finding motifs from highest attention tokens. Starting from the top 3 highest attention tokens, we first obtain the highest attention vocab for the positive and negative predicted samples, and from the token count vectors of the octamers in the vocab, we compute the mannwhitenyU p-values, and filtering for less than 0.01, we match the sequence motifs with the JASPAR TF motifs using FIMO, and return the corresponding p-values and q-values for those with pi0 FDR less than 0.5.

### 3.2 Ablation Studies

We found that pretraining using a dataset without offsets had a higher pretraining loss than while using a dataset with 3 offsets 1, 2 and 3. The validation dataset loss during pretraining reached 0.23 after 10000 steps when the dataset has 3 offsets, compared to a pretraining validation set loss of 0.52 using 1 offset and a pretraining validation set loss of 1.016 using no offset. Though the offsets had little to no impact during finetuning. During finetuning, the 0 offset method reported an accuracy of 79.05%, a precision of 78.82%, a recall of 76.93%, and an F1-score of 77.87%. The 1 offset model reported an accuracy of 77.95%, a precision of 77.01%, a recall of 76.93%, and F1-score of 76.97%. Moreover, the 3 offset model obtained an accuracy of 79.26%, a precision of 79.265%, a recall of 76.34%, and an F1-score of 77.77%. These results are summarized in Table 2. We see here that the model with 0 offsets reported the highest F1-score, whereas the model with 3 offsets reported the highest accuracy, whereas the model with 1 offset reported lower values of all the metrics, showing no correlation between the number of offsets and the validation set metrics during finetuning, even though the validation loss during pretraining showed an inverse correlation between number of offsets and validation loss. This demonstrates the ability of Byte Pair Encoding of finding the relevant tokens, even without providing the alternate words with different starting nucleotides, similar to the concept of *k*-mers.

**Table 2:** Comparison between metrics of llama + ReLoRA models trained on hg19 and validated on left out chromosome 1, using different number of offsets.

| Number of offsets | Accuracy | Precision | Recall | F1-score |
| --- | --- | --- | --- | --- |
| 0 offset | 79.05% | 78.82% | 76.93% | 77.87% |
| 1 offset | 77.95% | 77.01% | 76.93% | 76.97% |
| 3 offsets | 79.26% | 79.26% | 76.34% | 77.77% |

The llama-based model when trained using a bigger context window of 1024, leaving out chromosome 1 as the validation dataset, obtained an accuracy of 79.735%, a precision of 75.72%, a recall of 83.52%, and an F1 score of 79.425%. Whereas when trained using a smaller context window of 256, similarly leaving out chromosome 1 for validation, attained an accuracy of 75.5%, a precision of 76.74%, a recall of 71.89%, and an F1-score of 74.24%. While using a context window of 512, similarly leaving out chromosome 1 for validation, we obtained an accuracy of 79.26%, a precision of 79.26%, a recall of 76.34%, and an F1-score of 77.77%. This showed us that using a bigger context window consistently increases the performance metrics of our method, as can be seen in Table 3. We obtained the LOCOCV metrics for the human genome hg19 when trained using a context window of 1024 and obtained an average accuracy of 79.52±1.17%, an average precision of 79.03±2.04%, an average recall of 77.72±32%, and an average F1-score of 78.33±1.21%, as shown in Figure S4. This showed us that on an average, using a context length of 1024 provides us a 0.4% advantage in accuracy and a 0.2% advantage in F1-score than using a context length of 512. Trying out even larger context windows would require more GPU resources. The above results are compiled in Table 3

**Table 3:** Comparison between metrics of llama + ReLoRA models trained on hg19 and validated on left out chromosome 1, using different lengths of context windows.

| Context length | Accuracy | Precision | Recall | F1-score |
| --- | --- | --- | --- | --- |
| 256 | 75.50% | 76.74% | 71.89% | 74.24% |
| 512 | 79.26% | 79.26% | 76.34% | 77.77% |
| 1024 | 79.74% | 75.72% | 83.52% | 79.43% |

### 3.3 Comparisons

Upon training the transformer and bi-lstm based model on 24 chromosomes of the human genome hg19, it returned an average Leave One Chromosome Out Cross Validation (LOCOCV) accuracy of 74.43±1.51%, an average precision of 77.38±5.51%, and an average recall of 67.65±7.97%, along with an average F1 score of 71.62±3.37% for a context window of 512. For a context window of 1000, the model trained on hg19, returned a chromosome 1 accuracy of 72.83%, a precision of 72.58%, a recall of 68.15%, and an F1-score of 70.30%.

On the mouse genome mm9, the transformer and bi-lstm based model, attained a chromosome 1 accuracy of 71.88%, a precision of 83.87%, a recall of 50.04%, and an F1-score of 62.65%, using a context window of 1000 nucleotides.

Also, we have trained a transformer only model for capping prediction. On our transformer only model, the chromosome 1 accuracy on the hg19 genome using a context window of 1000 nucleotides, was found to be 72.24%, the precision was 77.86%, the recall was 57.52%, and the F1-score was at 66.12%. Using a context window of 500, our transformer model obtained a validation accuracy of 70.93%, a precision of 79.64%, a recall of 53.34%, and an F1 score of 63.89%.

The transformer and LSTM-based model, when trained using a neural network trained embedding layer instead of the alignment free contextual embeddings, yielded an accuracy of 29.42%, along with a precision of 27.42%, a recall of 29.38%, and an F1-score of 28.36%. However, modifying the input sequence to include 6-mers and completely removing the neural network trained embedding layer yielded an accuracy of 67.20%, a precision of 62.41%, a recall of 94.8%, and an F1-score of 75.27%.

We also trained an XGBoost based method from 64 element feature vectors, with the elements representing the frequency of the 3-mers found within the nucleotide sequence. The labels were kept the same as in TBCP. Using LOCOCV, the XGBoost yielded an accuracy of 74.49±1.31%, along with a precision of 76.86±3.05%, recall of 67.47±4.84% and F1-score of 73.68±1.89%. The LightGBM based model, trained using the same dataset, returned an LOCOCV accuracy of 74.35±1.18%, precision of 77.18±3.46%, recall of 66.69±5.58% and F1 score of 71.30±2.00%. The RF based model, also trained using the same dataset, returned an LOCOCV accuracy at 73.89±1.16%, precision of 77.70±3.85%, recall of 64.70±6.475% and F1 score of 70.27±2.37%.

The ML based models shown above have been compared in Figure S3. And the LOCOCV metrics for the llama and ReLoRa based model trained on the human genome hg19 using a different context window of 1024 is shown in Figure S4. Figure 4 provides a macro view of the metrics from the llama based and the machine learning based models. Table 4 summarizes the comparison metrics among the methods showcased above.

**Table 4:** Leave One Chromosome Out Cross Validation (LOCOCV) comparison of different Deep Learning methods used for capping classification in human genome hg19 (except for the last row, which is on the mouse genome mm9). Context length for all the methods are 512, except for the last two rows where it is **1024**. Transformer bi-LSTM uses freq. embeds.

| Method | Accuracy(%) | Precision(%) | Recall(%) | F1-score(%) |
| --- | --- | --- | --- | --- |
| Transformer bi-LSTM | 74.43±1.51 | 77.38±5.50 | 67.65±7.97 | 71.62±3.37 |
| Random Forest | 73.89±1.16 | 77.70±3.85 | 64.70±6.48 | 70.27±2.37 |
| LightGBM | 74.35±1.18 | 77.18±3.46 | 66.69±5.58 | 71.30±2.00 |
| XGBoost | 74.49±1.31 | 76.86±3.05 | 64.70±6.48 | 73.68±1.89 |
| Llama & ReLoRA based | 79.12±1.16 | 78.64±2.10 | 77.66±2.11 | 78.11±1.11 |
| **Llama & ReLoRA based** | 76.17±1.23 | 79.03±2.04 | 77.72±2.32 | 78.33±1.21 |
| **<u>Llama & ReLoRA based</u>** | 78.09±1.08 | 79.78±3.32 | 73.11±3.36 | 76.17±1.23 |

**Figure 4:**
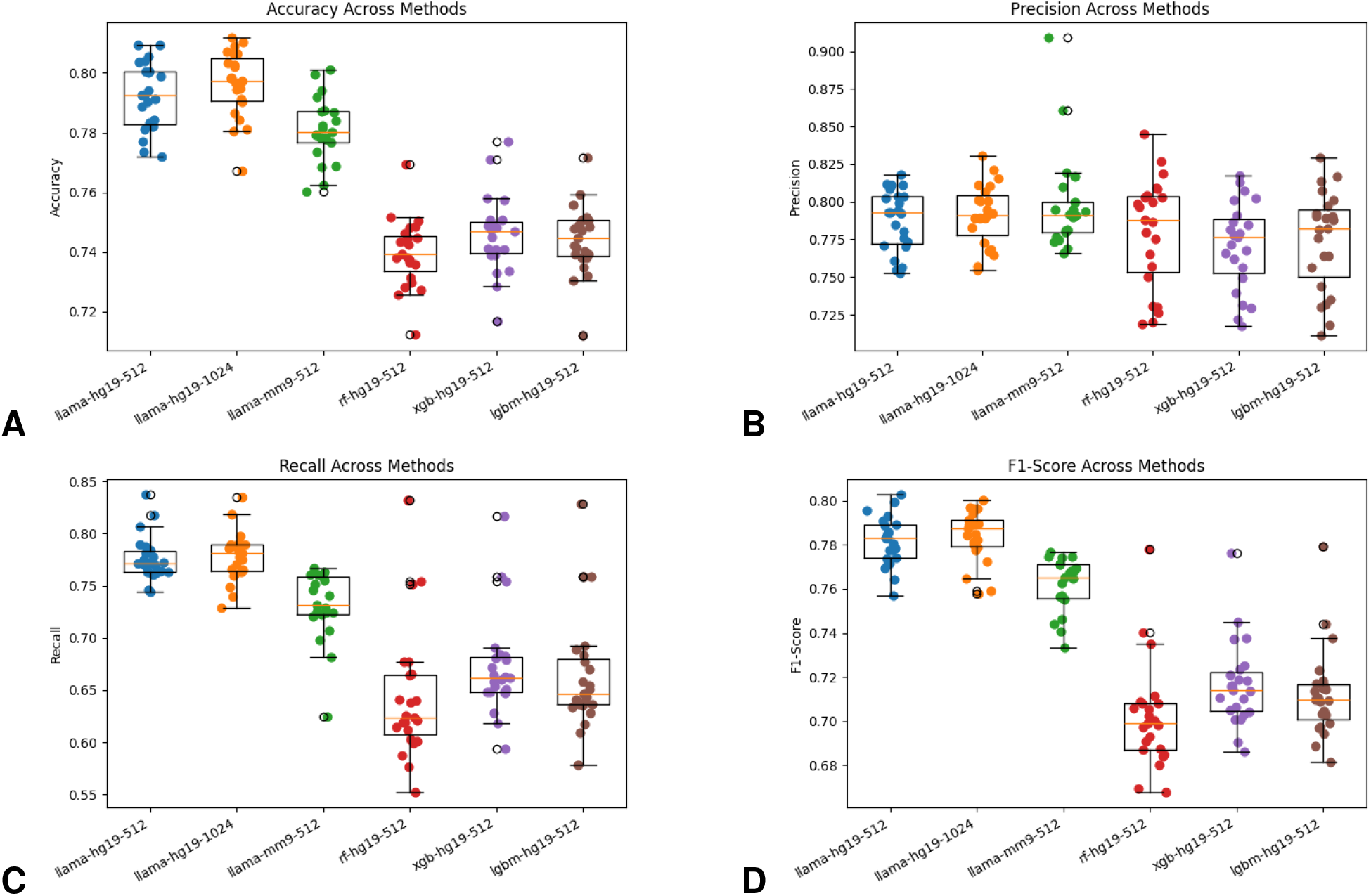
Box Plot comparison of A) accuracy B) Precision C) Recall and D) F1-score across all the main methods. Along the X-axis, we used the convention: MethodName-genome-contextWindowSize. Mehtod names are: Llama & ReLoRA based, Random Forest, XGBoost, and LightGBM. Genome is either human (hg19) or mouse (mm9) and the contenxt window is either 512 or 1024.

### 3.4 Hyperparameter Tuning

We trained each of the architectures through 5 epochs, optimized using the Adam optimizer with beta1 0.9, beta2 0.999, and epsilon 1e-08. We used a batch size of 64 and a linearly reducing learning rate of starting at 5e-5 for training the fine-tuned model. Using a lower batch size of 16 reported lower values in all validation. During validation on left out chromosome 1, a batch size of 64 reported an accuracy of 79.26% against an accuracy of 77.2% for batch size of 16, a precision of 79.26% against 76.645%, a recall of 76.34% against 75.365%, and an F1-score of 77.78% against 76.0% respectively. For pre-training, we had a batch size of 8, and used a learning rate scheduler, which first linearly increases the learning rate, but follows it up by reducing the learning rate on a sinosoidal curve, with downward spikes at fixed intervals.

## 4. Conclusion

CAGE sequencing has been used to measure the transcription expression levels of capped RNAs like mRNAs and lncRNAs. CAGE has been used as a longstanding tool to identify 5’ capping sites, but computational methods to predict capping sites has been scarce so far.

CAGE has been used in computational works to predict TSS, along with other methods like ChIPseq (Park, 2009) and ATACseq (Grandi et al., 2022) to filter out the noise from the signal. The source of noise being the capping sites that don’t coincide with promoter regions. There are capping sites in non-promoter regions like RNA splicing sites, which occurs when the RNA is recapped.(Berger et al., 2019; Kiss et al., 2015; A. Mukherjee et al., 2023) Some studies attribute approximately 25% of the total capping sites that lie in the downstream of TSS. So, predicting only the TSS site is not enough for mapping of all the capping sites. CAGE sites do not usually coincide to TSS signals like TATA boxes and initiators and the average distance of CAGE peaks to the TATA boxes and initiators were found to be 260 and 91 respectively. This necessitates the context window to be large enough to capture the TSS signal regions and the adjacent regions in order to predict capping. Our method takes a context window of 512 nts and predicts the occurrence of capping with an LOCOCV accuracy of 79.12±1.16%, a precision of 78.64±2.10%, a recall of 77.66±2.11%, along with an F1 score of 78.11±1.11%. On a larger context window of 1024, our method predicts capping with an LOCOCV average accuracy of 79.52±1.17%, an average precision of 79.03±2.04%, an average recall of 77.72±2.32%, and an average F1-score of 78.33±1.21%. Similarly, on fine-tuning on the mouse genome mm9, our method predicts the occurrence of capping with an LOCOCV accuracy of 78.09±1.08%, a precision of 79.78±3.32%, a recall of 73.105±3.36%, and an F1 score of 76.17±1.23%. It also returns regions of interest within the sequence having the maximum attention score. This region of interest has often shown co-occurrence to initiator signals and downstream promoter elements, with positive peaks mapping to initiators 0.133 times and to DPEs 0.578 times per positive case, and negative peaks mapping to initiators 0.231 times and to DPEs 0.590 times per negative case. Our model finds the capping regions in an input genomic sequence along with the tokens corresponding to the highest attention scores, from which we obtain the differentiating motifs for capping. These tokens have been found to be statistically significant when compared to the entire context length and against each other. Some of the prominent differentiating motifs found from the attention scores are TCTTGAAT, GTCTTGAA, ACTCATGT, CACTCATG, CCCT-CAGC, TGATCTGC, TTTTTATT, CAATACAT, GATGTGAG, TTGATCTG, AATA-CATA, ATTTAGGG, TAATTTCA, CTTGAATG, ATTCATAG, and ATGTGAGT, all of which have motif-wise p-values across predicted classes less than 0.01. Out of these motifs, TTTTTATT and ATGTGAGT are also biologically relevant since their resident sites match closely with some TF motifs.

## Data Availability

The entire code of this work is available at the following site: https://github.com/DibyoKgpIIT/CAGE_capping/tree/main.

## Acknowledgments

We acknowledge the use of the Paramshakti Supercomputer facility of the Indian Institute of Technology Kharagpur.

