## Supplementary Figures for "A Transformer based method for the Cap Analysis of Gene Expression and Gene Expression Tag associated 5’ cap site prediction in RNA"

*Correspondence to

Pralay Mitra, Ph.D.

Department of Computer Science and Engineering,

Indian Institute of Technology Kharagpur

West Bengal - 721302, India

ORCID ID: 0000-0003-4119-3788

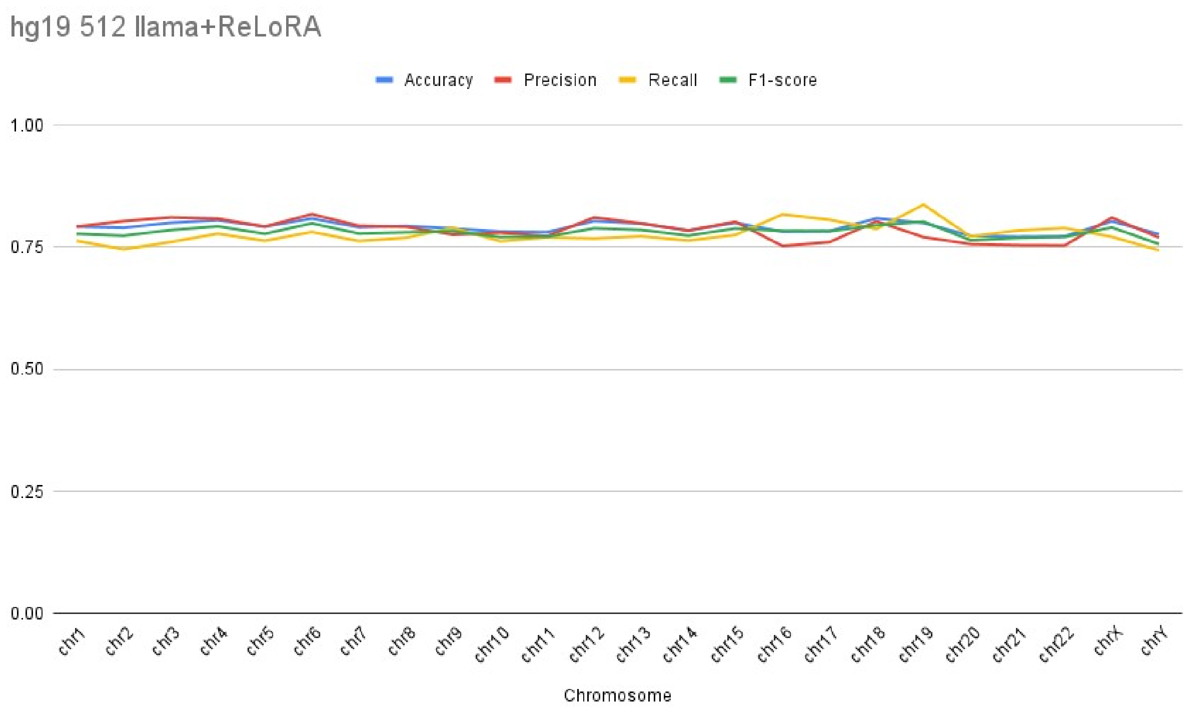


**Figure S1.** Plot showing the performance across metrics like accuracy, precision, recall, and F1-score validated on left out chromosomes 1 through 24 of the human genome hg19 using our Llama and ReLoRA based model.

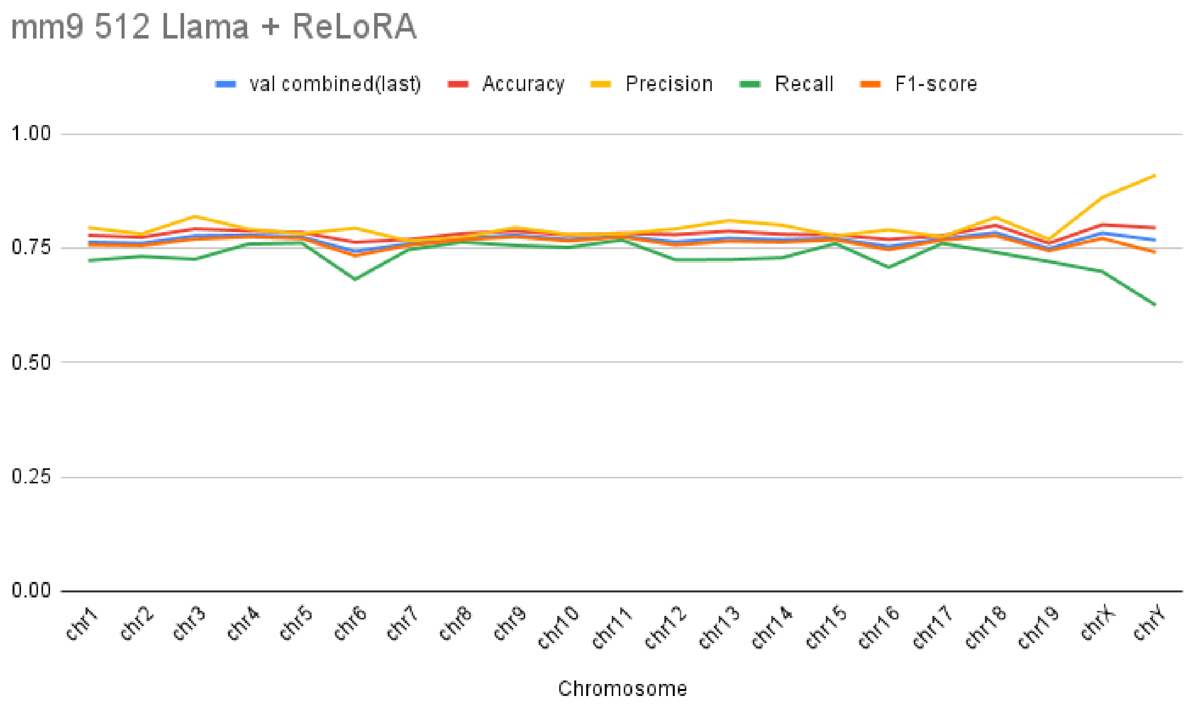


**Figure S2.** Plot showing the performance across metrics like accuracy, precision, recall, and F1-score validated on left out chromosomes 1 through 21 of the mouse genome mm9 using our Llama and ReLoRA based model.

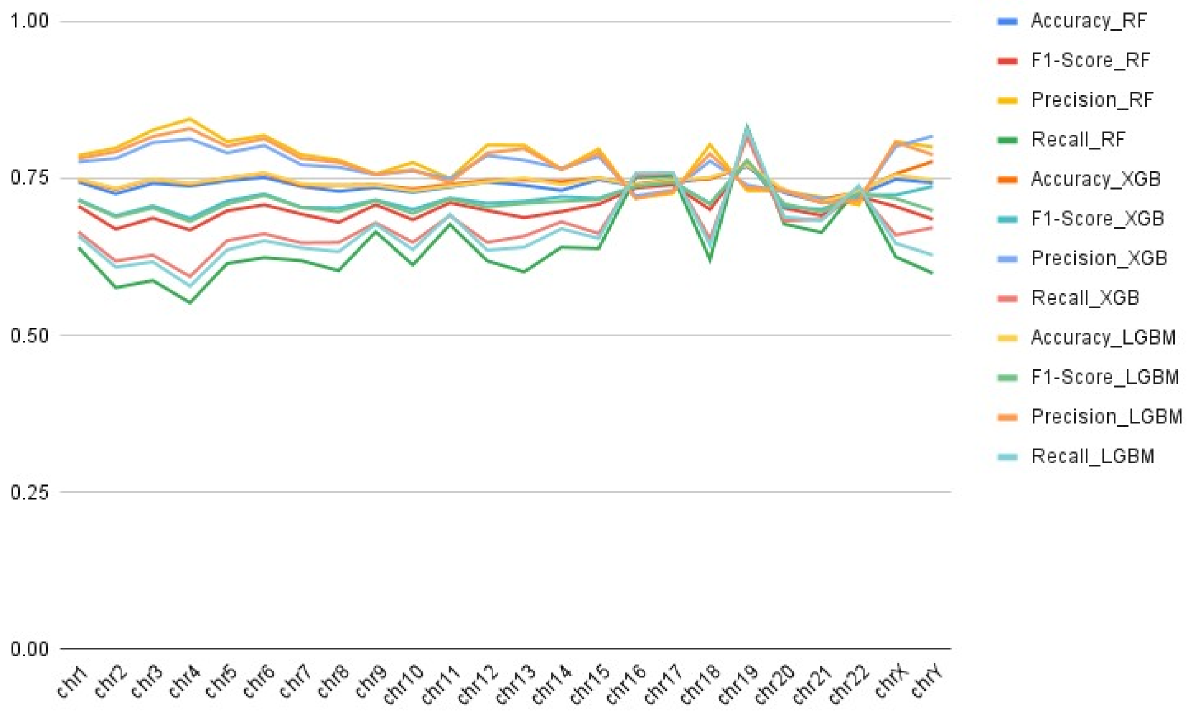


**Figure S3.** Comparison between Random Forest, XG Boost, and Light GBM in different metrics for LOCOCV on human genome hg19 using a context window of size 512.

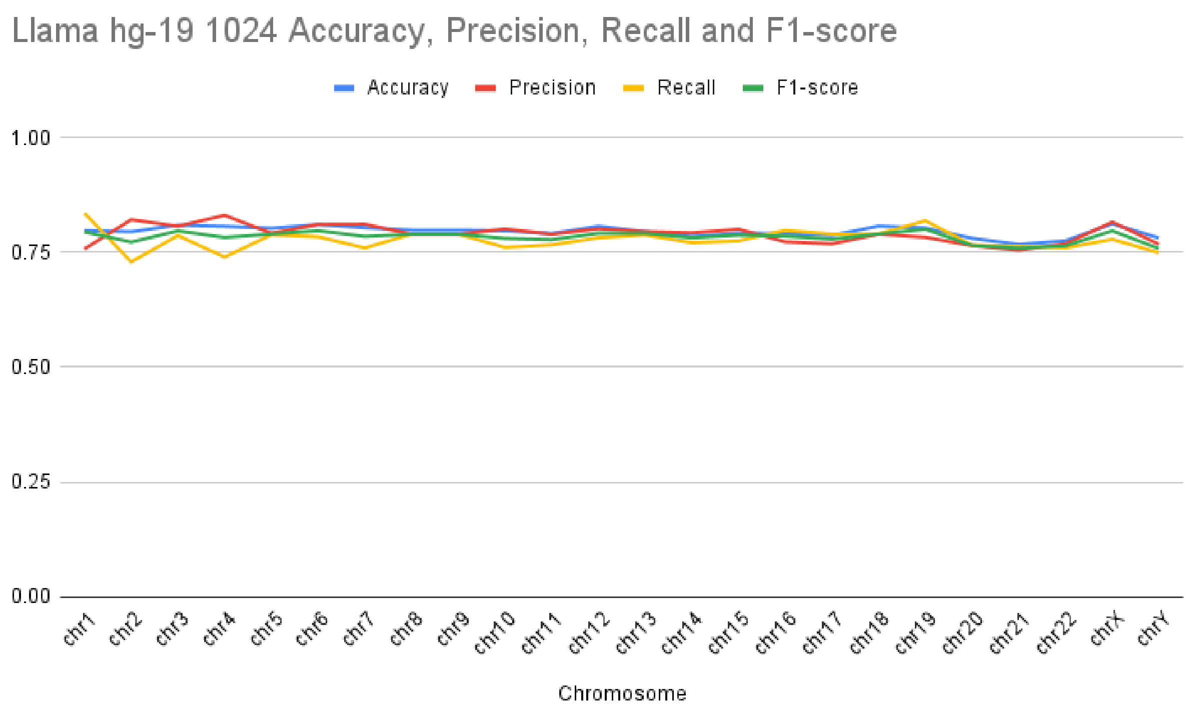


**Figure S4.** Llama 20 million model metrics for LCOCV on human genome hg19 for a context window of size 1024.
